# ATM orchestrates the DNA-damage response to counter toxic non-homologous end-joining at broken replication forks

**DOI:** 10.1101/330043

**Authors:** Gabriel Balmus, Domenic Pilger, Julia Coates, Mukerrem Demir, Matylda Sczaniecka-Clift, Ana Barros, Michael Woods, Beiyuan Fu, Fengtang Yang, Elisabeth Chen, Matthias Ostermaier, Tatjana Stankovic, Hannes Ponstingl, Mareike Herzog, Kosuke Yusa, Francisco Munoz Martinez, Stephen T. Durant, Yaron Galanty, Petra Beli, David J. Adams, Allan Bradley, Emmanouil Metzakopian, Josep V. Forment, Stephen P. Jackson

## Abstract

Mutations in the *ATM* tumor suppressor confer hypersensitivity to DNA-damaging agents. To explore genetic resistance mechanisms, we performed genome-wide CRISPR-Cas9 screens in cells treated with the DNA topoisomerase poison topotecan. Thus, we establish that loss of terminal components of the non-homologous end-joining (NHEJ) machinery or the BRCA1-A complex specifically confers topotecan resistance to ATM-deficient cells. We show that hypersensitivity of ATM-mutant cells to topotecan or the poly-(ADP-ribose) polymerase inhibitor olaparib is due to delayed homologous recombination repair at DNA-replication-fork-associated double-strand breaks (DSBs), resulting in toxic NHEJ-mediated chromosome fusions. Accordingly, restoring legitimate repair in ATM-deficient cells, either by preventing NHEJ DNA ligation or by enhancing DSB-resection by BRCA1-A complex inactivation, markedly suppresses this toxicity. Our work suggests opportunities for patient stratification in ATM-deficient cancers and when using ATM inhibitors in the clinic, and identifies additional therapeutic vulnerabilities that might be exploited when such cancers evolve drug resistance.

**One Sentence Summary:** ATM counteracts toxic NHEJ at broken replication forks

## INTRODUCTION

Recognition and repair of DNA damage is crucial for all organisms (Jackson and Bartek, 2009). DNA double-strand breaks (DSBs) are particularly toxic lesions, caused either directly by ionizing radiation (IR) and reactive chemicals, or indirectly by processing of other DNA lesions or breakdown of DNA replication forks. Upon detecting DSBs, cells activate the DNA damage response (DDR) signal-transduction pathway slowing or halting cell cycle progression to allow time for repair. DSB repair is mainly carried out by two complementary mechanisms, non-homologous end-joining (NHEJ) and homologous recombination repair (HRR) (Ciccia and Elledge, 2010).

Classical NHEJ is initiated by recruitment of the Ku70/80 heterodimer to DNA ends, which then engages with the DNA-dependent protein kinase catalytic subunit (DNA-PKcs) to form the DNA-PK complex that is subsequently stabilized on chromatin by PAXX. XRCC4, XLF and DNA ligase IV (LIG4) and recruited along with other factors to process, align and ligate the ends independently of sequence homology (Blackford and Jackson, 2017). HRR involves binding of the MRE11-RAD50-NBS1 (MRN) complex to tether DSB ends and recruit and activate the ataxia telangiectasia mutated (ATM) protein kinase (Blackford and Jackson, 2017). Together with CTIP, the MRN complex promotes resection of DSB ends to produce single-stranded DNA (ssDNA) overhangs that are protected by binding of replication protein A (RPA). RPA is then replaced by the recombinase RAD51, mediating strand invasion into the homologous sister chromatid to allow error-free repair (Huertas, 2010). RAD51 loading onto ssDNA requires the tumor suppressor proteins BRCA1 and BRCA2, deficiencies of which cause HRR defects and predisposition to breast, ovarian and other cancers (Nielsen et al., 2016). Although BRCA1 exists in several protein complexes (BRCA1-A, -B and -C), their contributions to the phenotypes of BRCA1-deficient cells are poorly understood (Wang, 2012).

While NHEJ is active in G0 and throughout interphase, HRR is restricted to S and G2 cell cycle phases, where a homologous sister chromatid is available as repair template. Several layers of control dictate DSB-repair-pathway choice between NHEJ and HRR, including activation of HRR by cyclin-dependent kinase activity (Chapman et al., 2012), or competition between HRR- and NHEJ-promoting factors (Ceccaldi et al., 2016). The latter involves regulating recruitment kinetics of the MRN and Ku complexes (Hustedt and Durocher, 2017), as well as MRN/CTIP-dependent removal of Ku from DSBs (Chanut et al., 2016a). While ATM modulates CTIP-dependent Ku removal (Chanut et al., 2016a), the impact of losing this function in ATM-deficient cells is unknown. Additionally, there is competition between HRR-promoting factor BRCA1 and NHEJ-promoting factor 53BP1, although the mechanisms underlying this antagonism are not clear (Panier and Boulton, 2014). The potential effects of other BRCA1 interacting proteins on DNA-end resection dynamics are not well studied, although defects in BRCA1-A complex components increase HRR efficiency in a manner linked to enhanced DSB resection (Prakash et al., 2015).

DSBs also arise when DNA replication-forks break down upon encountering DNA lesions such as single-strand breaks, or protein-DNA complexes such as abortive DNA topoisomerase I (TOP1) catalytic intermediates or inhibited/trapped poly(ADP-ribose) polymerase 1 (PARP1) enzyme when single-ended DSBs (seDSBs) are generated. These structures are not physiologically suited to NHEJ, instead being dealt with by break-induced replication (BIR) that uses the sister chromatid as template to restore the replication fork, a process that invariably produces sister chromatid exchanges (SCEs) (Helleday, 2003; 2011). Consequently, drugs producing seDSBs, such as the TOP1 poison camptothecin (CPT, and its derivatives topotecan and irinotecan) or PARP inhibitors such as olaparib, effectively kill HRR-defective cells. Indeed, selective killing of *BRCA1* or *BRCA2* mutant tumor cells by PARP inhibitors is now of established clinical utility, and may be extended to tumors with mutations in other genes, such as *ATM*, that are thought to share molecular features with *BRCA*-mutant cells (Lord and Ashworth, 2016).

Herein, we perform genome-wide CRISPR-Cas9 loss-of-function genetic screens to identify suppressors of cell killing by the TOP1 poison topotecan in ATM-proficient and ATM-deficient cells. Thus, in addition to highlighting potential mechanisms for therapeutic resistance in ATM-deficient cancers, our studies lead to a model in which the prime mechanism by which ATM promotes cell survival in response to seDSB generation is to promote efficient DSB resection to prevent seDSB repair by toxic NHEJ.

## RESULTS

### CRISPR-Cas9 screens for suppression of sensitivity to TOP1 poisons

We derived wild-type (WT) and *Atm*-null isogenic mouse embryonic stem cells (mESCs) from an ATM-deficient mice (Barlow et al., 1996). *Atm*^*-/-*^ mESCs exhibited complete loss of ATM protein and, upon DNA damage induction with CPT or IR, failed to mediate ATM-dependent signaling as measured by CHK2 Thr68 phosphorylation **(Fig. 1A)**. In line with previous reports (Choi et al., 2016), ATM-deficient mESCs were more sensitive than WT cells to the CPT derivative topotecan, which is used clinically to treat diverse cancers **(Fig. 1B,C)**.

**Figure 1.**
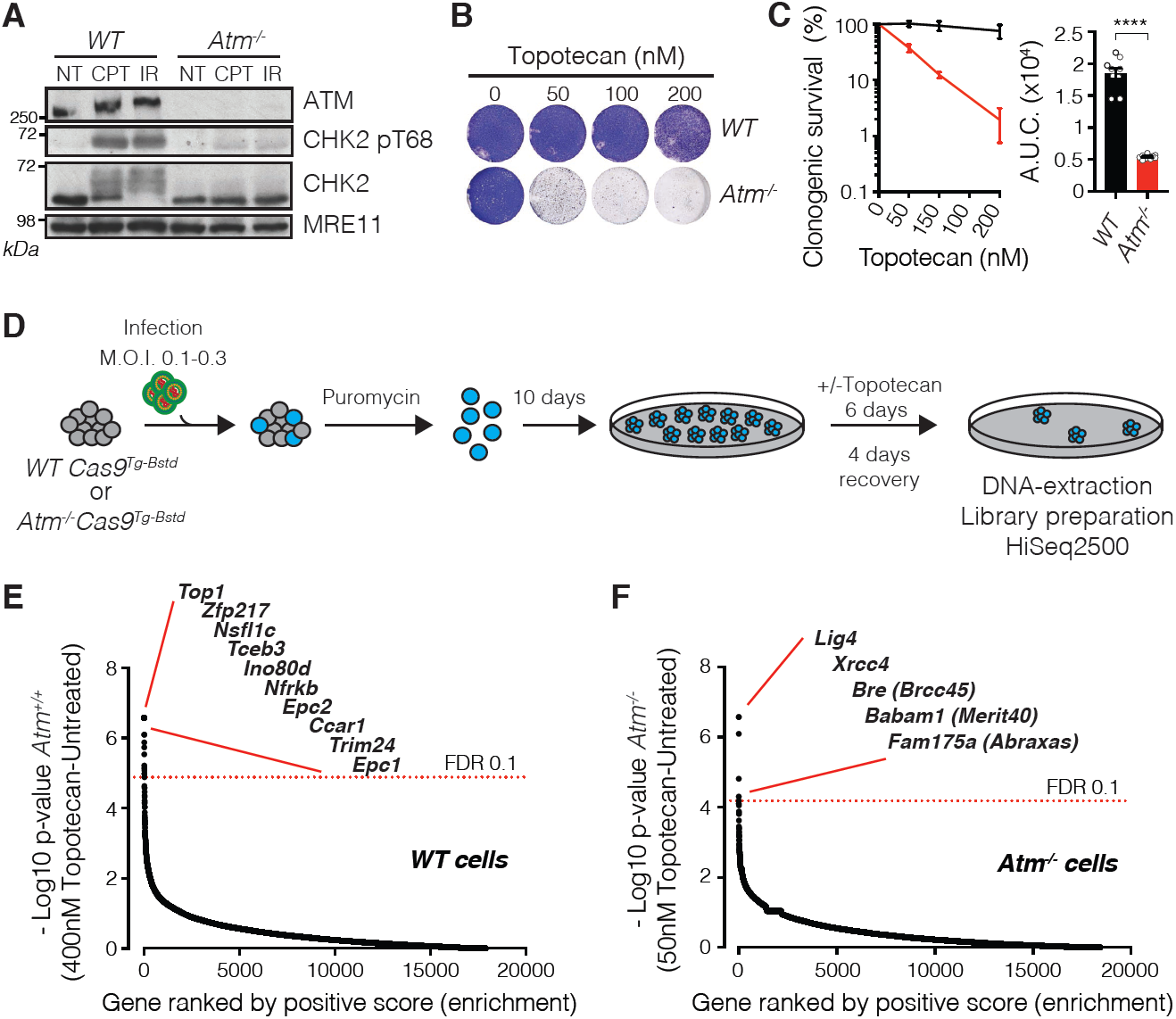
CRISPR-Cas9 screening in WT and ATM-deficient mESCs. **(A)** Representative western blot images show absence of ATM protein and defective signaling through phosphorylation of its substrate CHK2 on Thr-68. NT: untreated; CPT: camptothecin (1µM, 1h); IR: ionizing radiation (10Gy, 1h). MRE11 is used as loading control. **(B, C)** Crystal violet cell viability assay **(B)** and clonogenic survival assays **(C)** showing hypersensitivity of ATM-deficient cells to topotecan; n=9/genotype; error bars SEM; t=15.17; df=4; ****p<0.0001; two-tailed Student’s t test based on AUC (Area Under the Curve). **(D)** Outline of the CRISPR screen. Wild-type (WT) or ATM-deficient cells stably expressing Cas9 nuclease were infected with lentiviral particles containing the whole-genome sgRNA library, subjected to puromycin selection, and passaged to ensure loss of affected protein products. Puromycin-resistant WT or *Atm*^*-/-*^ cells were exposed, respectively, to 400nM and 50nM topotecan for 6 days, and resistant pools isolated. Genomic DNA was extracted from these and from parallel cell cultures treated in the absence of topotecan, and DNA libraries were prepared and sequenced using HiSeq2500. M.O.I: multiplicity of infection. **(E, F)** Classification of the most enriched CRISPR-targeted genes in topotecan-resistant WT **(E)** and *Atm*^*-/-*^ **(F)** mESCs. Dotted red lines represent positive enrichment false-discovery-rate (FDR) thresholds. Represented are the names of top hits with highest enrichment scores. All data were analyzed by using MAGeCK and are available in **Datasets S1**; **S2** Panels containing clonogenic survival assays (left) and area-under-curve (AUC; right) were generated using GraphPad Prism 7. Bars represent mean ± SEM; ****p<0.0001; ***p<0.001; **p<0.01; *p<0.05; ns= not significant (p>0.05); two-tailed Student’s t test following F test to confirm equal variance; df=4. For each clonogenic experiment data is pooled from n=3 individual experiments.

To explore mechanisms of topotecan resistance, we expanded WT or *Atm*^*-/-*^ clones expressing Cas9 nuclease **(Fig. S1A**), and transduced them with a pooled genome-wide lentiviral CRISPR small-guide RNA (sgRNA) library (Koike-Yusa et al., 2014). We treated the WT and *Atm*^*-/-*^ mESC cells with concentrations of topotecan pre-determined to give >90% cell-killing (**Fig. 1D; Fig. S1B**). Surviving cell pools were isolated, and the regions encoding sgRNAs PCR-amplified and subjected to next-generation DNA sequencing. sgRNA counts from topotecan-treated cells were compared with those from untreated cells. As expected, the most enriched sgRNAs in topotecan-selected WT cells corresponded to TOP1, the drug target (Pommier, 2006) **(Fig. 1E; Fig. S1C; Datasets S1 and S2)**. Amongst the other enriched sgRNA’s were those corresponding to genes encoding the TRRAP, EPC1, EPC2 and ING3 subunits of the NuA4 chromatin-remodeling complex (House et al., 2014) **(Fig. 1E)**, suggesting that this complex may promote topotecan toxicity in WT cells.

In contrast, in topotecan-resistant *Atm*^*-/-*^ cells, the most enriched sgRNAs targeted genes encoding core NHEJ factors XRCC4 and LIG4, or the BRCA1-A components BRCC45 (BRE), ABRAXAS (FAM175A) and MERIT40 (BABAM1) **(Fig. 1F; Fig. S1D;Datasets S3 and S4)**. However, sgRNAs targeting *Brca1* or genes for factors present in other BRCA1-containing complexes were not enriched (**Dataset S3)**. While interesting to examine many of these factors for their impacts on seDSB generation and repair and/or on associated cellular responses, for our ensuing studies we focused on NHEJ and BRCA1-A components in the context of ATM deficiency.

### NHEJ and BRCA1-A components mediate topotecan toxicity in ATM-null cells

To validate impacts of BRCA1-A components on the topotecan sensitivity of ATM-deficient cells, we used CRISPR-Cas9 gene editing to generate *Atm*^*-/-*^*Brcc45*^*-/-*^, *Atm*^*-/-*^ *Merit40*^*-/-*^, *Atm*^*-/-*^*Abraxas*^*-/-*^ and *Atm*^*-/-*^*Brcc36*^*-/-*^ double mutant mESCs (**Fig. 2A**; as found previously (Bin Wang et al., 2009; Feng et al., 2009; Shao et al., 2009), some BRCA1-A components were destabilized in the absence of certain other components). Absence of each of these genes suppressed topotecan toxicity in ATM-deficient mESCs (**Fig. 2B**), thus validating results from the screen. Similarly, we generated *Atm*^*-/-*^*Lig4*^*-/-*^ and *Atm*^*-/-*^*Xrcc4*^*-/-*^ double mutant cells by gene editing (**Fig. 2C**; as shown previously (Sibanda et al., 2001), loss of XRCC4 led to LIG4 destabilization). Strikingly, inactivation of *Xrcc4* or *Lig4* in *Atm*^*-/-*^ mESCs made them almost as resistant to topotecan as WT cells (**Fig. 2D, E**). Furthermore, re-expression of XRCC4 in *Atm*^*-/-*^ *Xrcc4*^*-/-*^ cells restored topotecan hypersensitivity (**Fig. S2A**). Importantly, the effects of XRCC4 or LIG4 loss on topotecan resistance were specific to *Atm*^*-/-*^ cells, as inactivating *Xrcc4* or *Lig4* in ATM-proficient cells (**Fig. S2B**) did not visibly enhance topotecan resistance (**Fig. S3C**) but did confer IR hypersensitivity (**Fig. S2D**). In stark contrast, combined loss of ATM and either XRCC4 or LIG4 caused cells to be more sensitive to IR than cells lacking ATM alone (**Fig. 2F**). These findings may reflect ATM and NHEJ playing complementary roles in responding to IR-induced two-ended DSBs, while acting in antagonistic ways at seDSBs arising during DNA replication.

**Figure 2.**
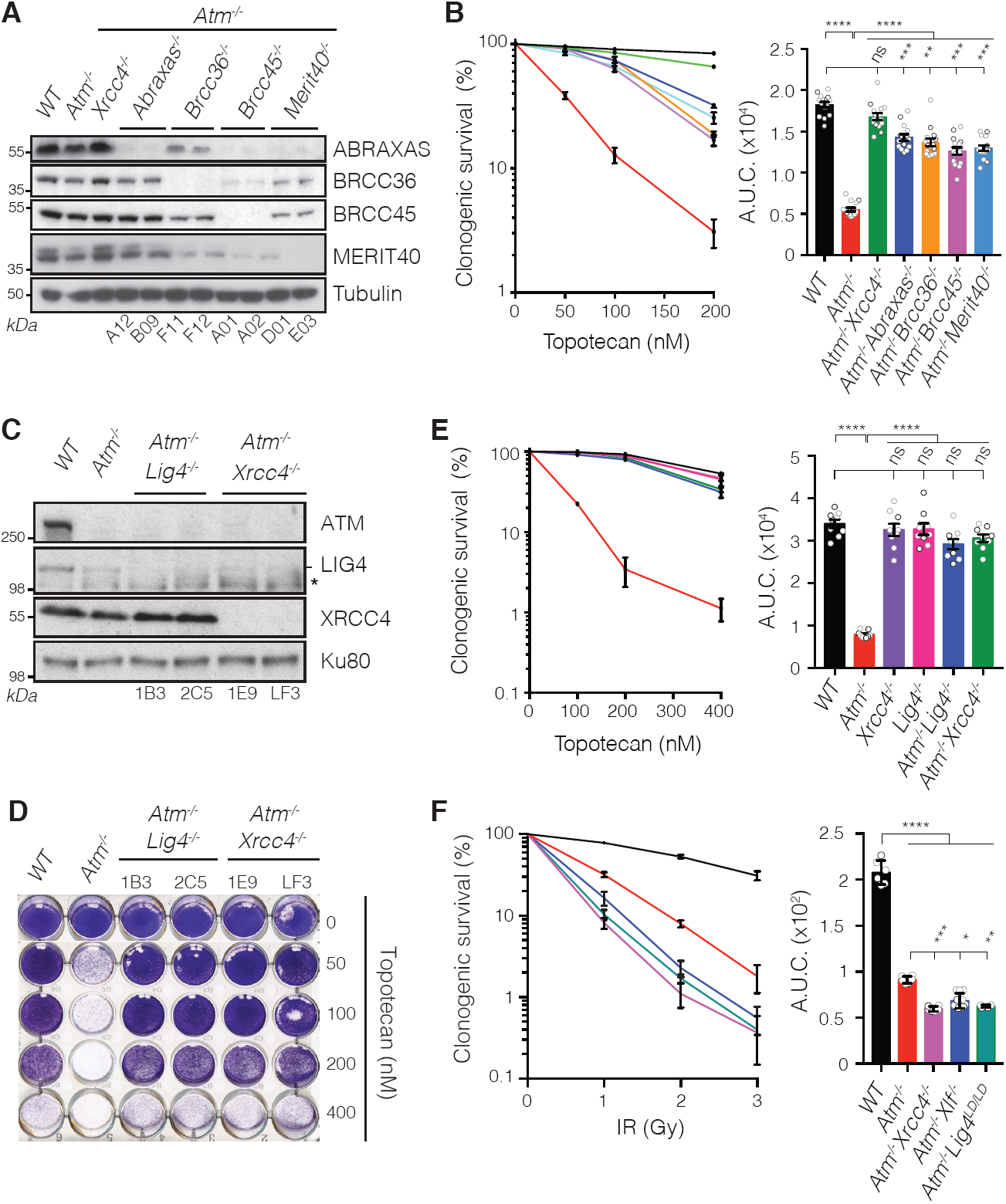
Loss of NHEJ factors or BRCA1-A complex members confers resistance to topotecan in ATM-deficient cells. **(A)** Representative western blot images depicting abundance of ABRAXAS, BRCC36, BRCC45 and MERIT40 proteins in *Atm*^*-/-*^*Abraxas*^*-/-*^, *Atm*^*-/-*^*Brcc36*^*-/-*^, *Atm*^*-/-*^*Brcc45*^*-/-*^ and *Atm*^*-/-*^*Merit40*^*-/-*^ cells as compared to *WT*, *Atm*^*-/-*^ and *Atm*^*-/-*^*Xrcc4*^*-/-*^cells. Tubulin is used as loading control. Two independent clones (clone numbers at the bottom) were used per genotype. **(B)** Quantification of clonogenic survival after topotecan treatment in *Atm*^*-/-*^*Abraxas*^*-/-*^ (n=15), *Atm*^*-/-*^*Brcc36*^*-/-*^ (n=15), *Atm*^*-/-*^*Brcc45*^*-/-*^ (n=15), *Atm*^*-/-*^*Merit40*^*-/-*^ (n=15) and *Atm*^*-/-*^ *Xrcc4*^*-/-*^ (n=15) cells as compared to *WT* (n=15) and *Atm*^*-/-*^ (n=15) cells. (**C**) Representative western blot images depicting abundance of ATM, LIG4 (-indicates LIG4; * indicates an antibody cross-reacting protein) and XRCC4 proteins in *Atm*^*-/-*^ *Lig4*^*-/-*^ and *Atm*^*-/-*^*Xrcc4*^*-/-*^ cells as compared to *WT* and *Atm*^*-/-*^ cells. Ku80 is used as loading control. **(D, E)** Crystal violet cell viability assay **(D)** and quantification of clonogenic survival **(E)** indicating suppression of *Atm*^*-/-*^ (n=9) dependent hypersensitivity to topotecan in *Atm*^*-/-*^*Lig4*^*-/-*^ (n=9) and *Atm*^*-/-*^*Xrcc4*^*-/-*^ (n=9) cells as compared to *WT* cells (n=9). Note that *Lig4*^*-/-*^ (n=9) or *Xrcc4*^*-/-*^ (n=9) single mutants do not exhibit increased topotecan resistance. **(F)** Quantification of clonogenic survival showing that *Atm*^*-/-*^*Xrcc4*^*-/-*^ (n=6), *Atm*^*-/-*^*Xlf*^*-/-*^ (n=10) *and Atm*^*-/-*^*Lig4*^*LD/LD*^ (n=6) cells are more sensitive to IR than *Atm*^*-/-*^ (n=6) cells. All clonogenic survival curves (left) and AUCs (right) were generated by using GraphPad Prism 7. Bars represent mean ± SEM; ****p<0.0001; ***p<0.001; **p<0.01;*p<0.05; ns= not significant (p>0.05); two-tailed Student’s t test following F test to confirm equal variance; df=4 (3 independent experiments; n=5 in panel (**B**); n=3 in panel (**E**); n=2 in panel (**F**) for each experiment).

### Topotecan toxicity in ATM-deficient cells is mediated by LIG4 catalytic activity

To complement our mESC studies, we generated and validated *ATM*^*-/-*^, *LIG4*^*-/-*^ *and ATM*^*-/-*^*LIG4*^*-/-*^ clones in human RPE1 cells (**Fig. S2E-I**). *ATM*^*-/-*^*LIG4*^*-/-*^ cells were more resistant to CPT than *ATM*^*-/-*^ cells (**Fig. S2J**). Furthermore, chemical inhibition of ATM kinase activity (ATMi) (Hickson et al., 2004) in WT RPE1 cells sensitized them to topotecan, while LIG4-deficiency partially suppressed this phenotype (**Fig. S2K**). These results thus indicated that ATM kinase activity prevents CPT-induced cell killing by a mechanism that relies, at least in part, on LIG4.

Absence of XRCC4 resulted in decreased LIG4 protein levels, but not vice versa (**Fig. 2C**), suggesting that LIG4 mediates sensitivity of ATM-null cells to TOP1 poisons. To evaluate whether LIG4 DNA ligase activity was required, we generated the K273A point mutation that abrogates LIG4 catalytic function (Cottarel et al., 2013) in *Atm*^*-/-*^ mESCs (**Fig. S3A, B**). Similar to complete loss of LIG4, catalytically inactive *Lig4*^*LD/LD*^ conferred strong resistance to topotecan (**Fig. 3A, B**) but not IR (**Fig. 2F**) in *Atm*^*-/-*^ cells. These observations implicated DNA ligation activity, rather than a structural function of the XRCC4-LIG4 complex, as mediating topotecan toxicity in the absence of ATM. To extend our findings to a more physiological setting, we generated mouse tumor xenografts using our mESC lines, and treated the mice with topotecan (Pawlik et al., 1998). In agreement with our *in vitro* data, *Atm*^*-/-*^ tumors were highly sensitive to topotecan when compared to WT controls, while *Atm*^*-/-*^*Xrcc4*^*-/-*^ tumors showed increased drug resistance (**Fig. 3C**). Collectively, these findings highlighted that LIG4 catalytic activity is a major driver for topotecan toxicity in cells lacking functional ATM, and establish that LIG4-XRCC4 function confers topotecan hypersensitivity to ATM-deficient cells both *in vitro* and *in vivo*.

**Figure 3.**
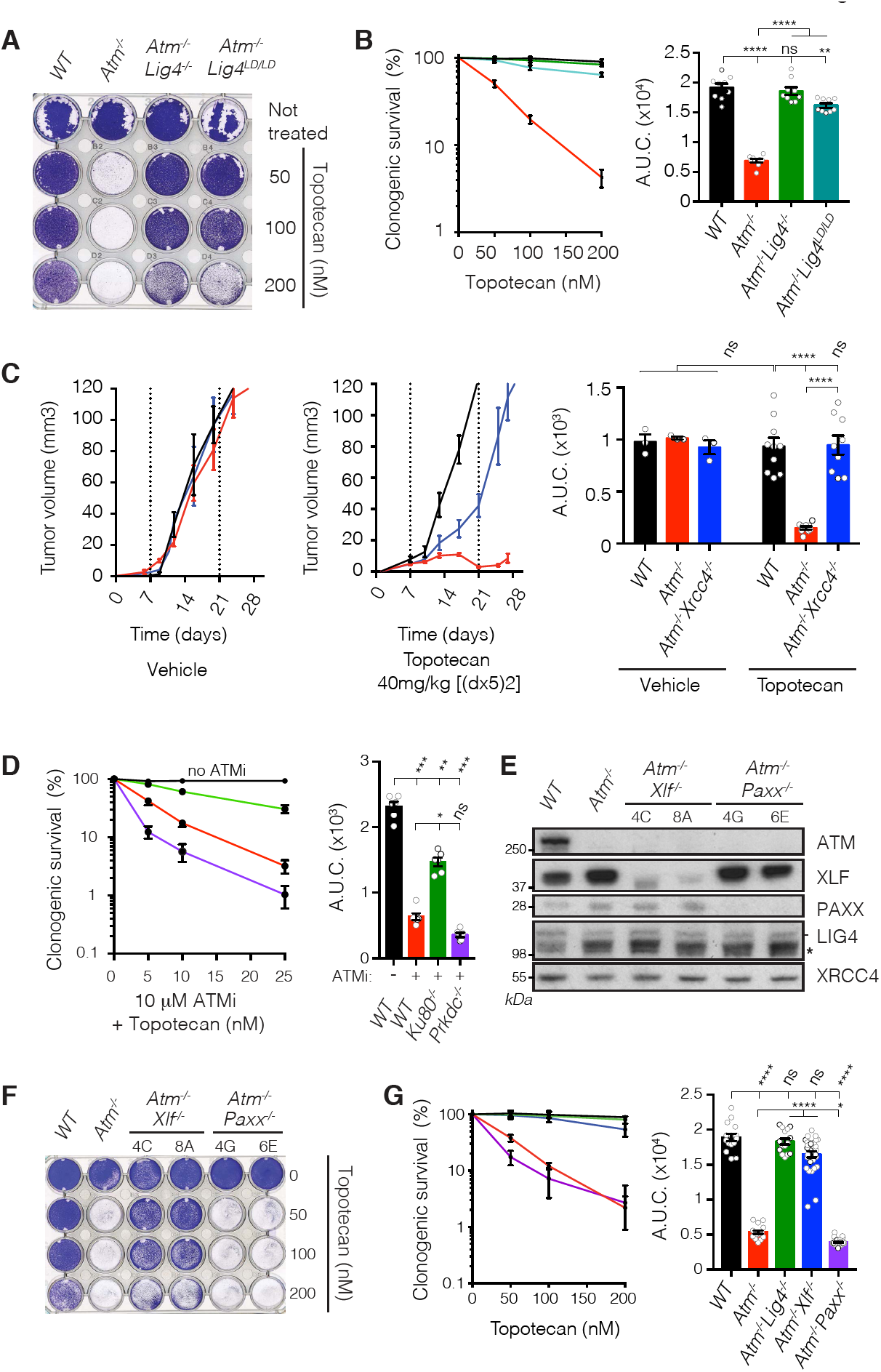
LIG4 catalytic activity mediates topotecan sensitivity in ATM-deficient cells. **(A)** Crystal violet cell viability assay shows that LIG4 catalytic activity mediates hypersensitivity of ATM-deficient cells to topotecan. LD: ligase-dead allele. **(B)** Quantification of clonogenic survival indicating suppression of *Atm*^*-/-*^ (n=9) cell hypersensitivity to topotecan upon abrogation of LIG4 catalytic activity in *Atm*^*-/-*^ *Lig4*^*LD/LD*^ (n=9) as compared to WT (n=9) and *Atm*^*-/-*^*Lig4*^*-/-*^ (n=9). Data from n=3 individual experiments. **(C)** Mouse xenograft studies indicating that *Atm*^*-/-*^*Xrcc4*^*-/-*^ deficient tumors are more resistant than *Atm*^*-/-*^ single mutant tumors to 40mg/kg topotecan treatment (days 1-5 and 8-12 equivalent to [(dx5)2] schedule *via* i.p. injections). Growth of untreated tumors (n=3 mice/genotype) was compared to growth of topotecan treated tumors (n=9 mice/genotype) (left) and AUC calculated and graphed (right). **(D)** Quantification of clonogenic survival assays showing that inhibiting ATM kinase activity sensitizes WT cells to topotecan and that inactivation of *Xrcc5/Ku80* but not *Prkdc/DNA-PKcs* partially suppresses this phenotype. n=9/genotype. (**E**) Representative western blot images depicting abundance of ATM, XLF and PAXX proteins in *Atm*^*-/-*^*Xlf*^*-/-*^ and *Atm*^*-/-*^*Paxx*^*-/-*^ cells as compared to *WT* and *Atm*^*-/-*^ cells. LIG4 (-indicates LIG4; * indicates an antibody cross-reacting protein) and XRCC4 used as loading controls. **(F)** Crystal violet cell viability assay showing that loss of XLF but not of PAXX ameliorates hypersensitivity of ATM-deficient cells to topotecan. **(G)** Quantification of clonogenic survival indicating that hypersensitivity of *Atm*^*-/-*^ cells to topotecan is alleviated in *Atm*^*-/-*^*Xlf*^*-/-*^ but not in *Atm*^*-/-*^*Paxx*^*-/-*^ cells (WT and *Atm*^*-/-*^*Lig4*^*-/-*^ cells used as controls). n=15/genotype. For the panel containing clonogenic survival assays as well as tumor volume percentage survival curves (left) and AUC (right) were generated by using GraphPad Prism 7. Bars represent mean ± SEM; ****p<0.0001; ***p<0.001; **p<0.01; *p<0.05; ns= not significant (p>0.05); two-tailed Student’s t test following F test to confirm equal variance; df=4 in panel (**B**), df=12 for the untreated and df=16 for the topotecan treated mice in panel (**C**) and df=4 for panels **(D)** and **(G)**. For panel **(E)**, individual clone names are represented below the genotypes. Data from n=3 individual experiments.

### Only some NHEJ factors mediate topotecan sensitivity in ATM-null cells

Another core NHEJ component is the DNA-PK complex, comprising the Ku70/80 heterodimer and DNA-PKcs (Blackford and Jackson, 2017; Gottlieb and Jackson, 1993). As combined genetic inactivation of ATM and DNA-PK is lethal to mouse cells (Sekiguchi et al., 2001), we generated *Prkdc* (encoding DNA-PKcs) or *Xrcc6* (encoding Ku80) deficient mESCs (**Fig. S3C-F**), then treated these and WT cells with a combination of ATMi plus various concentrations of topotecan. In contrast to the effect of Ku80 loss ((Britton et al., 2013), DNA-PKcs-deficiency did not increase topotecan resistance in the context of ATM inhibition (**Fig. 3D**). We also generated *Atm*^*-/-*^*Xlf*^*-/-*^ and *Atm*^*-/-*^*Paxx*^*-/-*^ cells (**Fig. 3E**), as XLF and PAXX play partially redundant roles in NHEJ (Balmus et al., 2016; Kumar et al., 2016; Lescale et al., 2016; Liu et al., 2017; Tadi et al., 2016). This revealed that absence of XLF, but not PAXX, significantly suppressed topotecan hypersensitivity in *Atm*^*-/-*^ cells (**Fig. 3F, G**). Collectively, our data indicated that hypersensitivity of ATM-deficient cells to TOP1 poisoning is mediated by toxic reactions arising from a subset of NHEJ components, likely *via* them promoting LIG4 catalytic activity towards seDSBs arising during DNA replication.

### Resistance to combined TOP1 poisoning and ATM inhibition in cancer cells

Combination treatment with ATMi and TOP1 poisons has been proposed to induce synergistic killing of cancer cells (Pommier, 2006). Indeed, combination of the TOP1 poison SN-38 and the ATMi AZD0156 is currently being evaluated clinically (NCT02588105), with special emphasis in colorectal cancer. To assess whether the resistance mechanisms we identified might be relevant in this setting, we conducted a CRISPR-Cas9 screen in the colorectal cancer cell line HT-29 (**Fig. 4A; Dataset S5**). Strikingly, the top gene hits that suppressed sensitivity to the combined action of SN-38 and AZD0156 encoded proteins of the NHEJ and BRCA1-A complexes (**Fig. 4B**; for additional hits see **Dataset S5**). This indicates conservation of the suppression mechanism in a human cancer cell line and that it not only operates when ATM protein is absent, but also when its catalytic activity is inhibited. Taken together, these data suggest that exploring the genetic status of NHEJ and BRCA1-A components in ATM-deficient tumors, or when exploring drug combinations with ATMi, might help identify which patients are most likely to benefit from agents such as TOP1 inhibitors.

**Figure 4.**
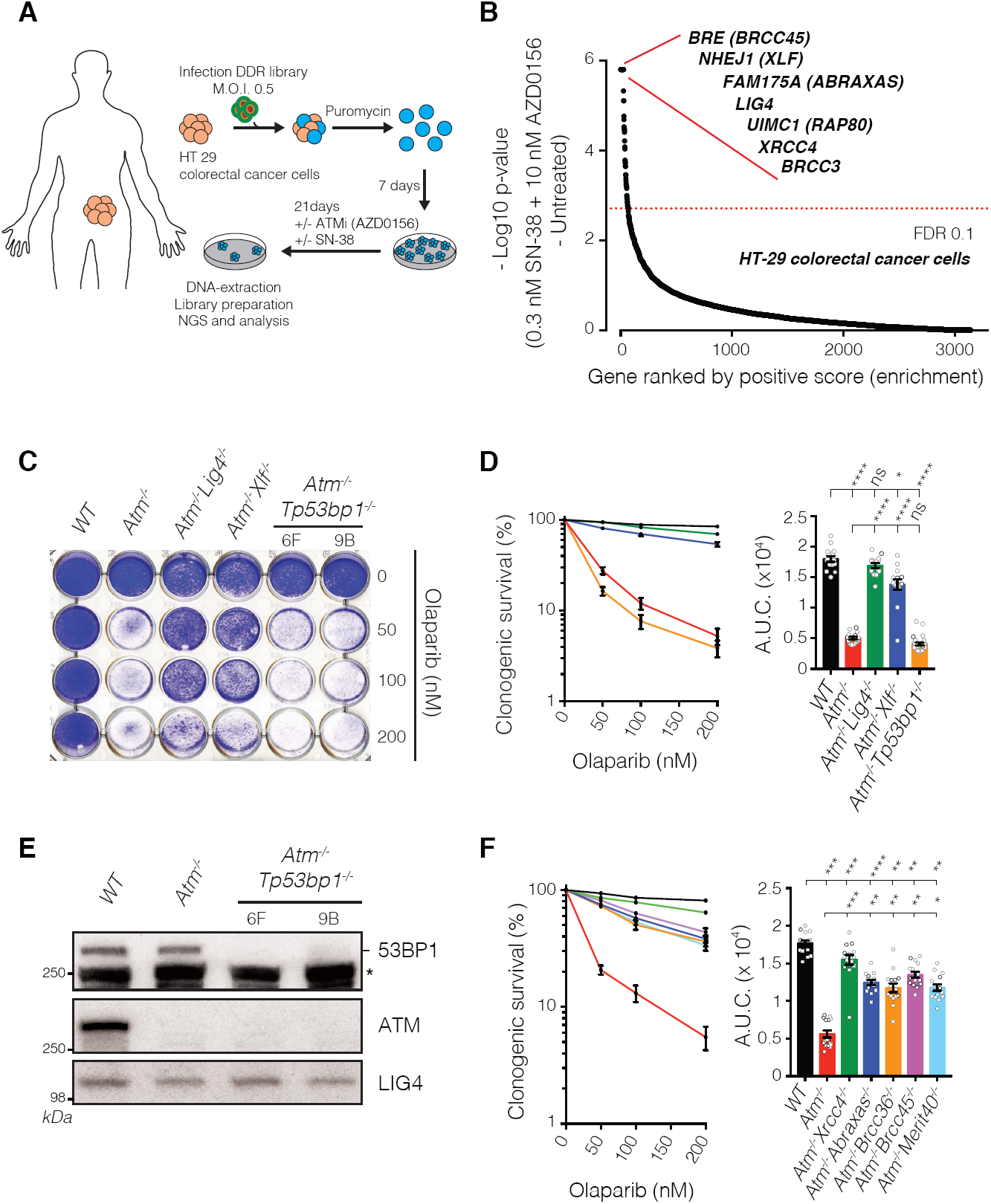
Mechanism of suppression in ATM-deficient cells is different to that in BRCA1-deficient cells. **(A)** Outline of the CRISPR screen in cancer cells. HT-29 colorectal cancer cells were infected with lentiviral particles containing the whole-genome sgRNA library, subjected to puromycin selection, and passaged to ensure loss of affected protein products. Puromycin-resistant cells were exposed to 10nM ATMi (AZD0156) and 0.3nM Irinotecan (SN-38) for 21 days, and resistant pools were isolated. Genomic DNA was extracted from these and from parallel cell cultures treated in the absence of topotecan, and DNA libraries were prepared and sequenced. M.O.I: multiplicity of infection. **(B)** Classification of the most enriched CRISPR-targeted genes. Dotted red lines represent positive enrichment false-discovery-rate (FDR) thresholds. Represented are the names of top hits with highest enrichment scores. All data were analyzed by using MAGeCK and are available in **Dataset S5. (C)** Crystal violet cell viability assay showing that *Atm*-mutant mESCs are hypersensitive to the PARP inhibitor olaparib. Inactivation of *Lig4* or *Xlf*, but not of *Tp53bp1*, rescues the olaparib hypersensitivity of *Atm*-deficient cells. **(D)** Quantification of clonogenic survival showing that loss of XLF (n=15) but not 53BP1 (n=30) can suppress the hypersensitivity of *Atm*^*-/-*^ cells (n=15) to olaparib as compared to WT control (n=15).**(E)** Representative western blot images depicting 53BP1 (-indicates 53BP1; * indicates antibody cross-reacting proteins) and ATM protein levels in *Atm*^*-/-*^*Trp53bp1*^*-*^*/-* cells as compared to WT and *Atm*^*-/-*^ cells. LIG4 used as loading control. **(F)** Quantification of clonogenic survival showing significant rescue of *Atm*^*-/-*^ dependent sensitivity to olaparib upon loss of individual BRCA1-A complex members in *Atm*^*-/-*^ *Abraxas*^*-/-*^, *Atm*^*-/-*^*Brcc36*^*-/-*^, *Atm*^*-/-*^*Brcc45*^*-/-*^ and *Atm*^*-/-*^*Merit40*^*-/-*^ cells. n=15/genotype. Panels **(D)** and **(F)** containing clonogenic survival curves (left) and AUC (right) were generated using GraphPad Prism 7. Bars represent mean ± SEM; ****p<0.0001; ***p<0.001; **p<0.01; *p<0.05; ns= not significant (p>0.05); two-tailed Student’s t test following F test to confirm equal variance. df=4 **(D)** and df=4 **(F)**. Data from n=3 individual experiments. For panel **(C)**, individual clone names are represented below the genotypes.

### 53BP1 loss does not suppress topotecan or olaparib hypersensitivity of ATM-deficient cells

Similar to TOP1 poisons, PARP1 inhibitors cause replication-fork breakage and seDSBs that require HRR (Helleday, 2011). In line with this and our findings with topotecan and CPT, *Atm*^*-/-*^ cells displayed hypersensitivity to olaparib that was suppressed by inactivating *Lig4* or *Xlf* (**Fig. 4C, D**). This suggested that a similar NHEJ-mediated toxicity mechanism operates for both topotecan and olaparib in ATM-deficient cells.

In experimental and therapeutic contexts, inhibitors of TOP1 or PARP1 are particularly toxic to cells with mutations in the HRR genes *BRCA1* and *BRCA2*. Furthermore, there is growing data indicating similar toxicities in cells with mutations in genes such as *ATM* that are thought to share molecular features with *BRCA*-mutant cells (Lord and Ashworth, 2016). Indeed, terms such as “BRCA-ness” or “HR-deficiency” (HRD) are used to highlight functional similarities between such genetic defects. Intriguingly, our genome-wide screen for topotecan resistance in ATM-null cells did not identify *Tp53bp1*, a gene whose inactivation restores HRR proficiency to *BRCA1*-mutant cells and confers PARP-inhibitor resistance (Bouwman et al., 2010; Bunting et al., 2010). In accord with our screening data, unlike loss of LIG4, XRCC4 or XLF, 53BP1 inactivation (**Fig. 4E**) did not suppress the hypersensitivity of ATM-deficient cells towards olaparib or topotecan **(Fig. 4C, D; Fig. S3G)**. Furthermore, in contrast to 53BP1 loss, genetic inactivation of BRCA1-A components rescued olaparib hypersensitivity in ATM-deficient cells (**Fig. 4F**). These data thus suggested that hypersensitivity to olaparib in ATM-deficient cells is mechanistically different from that in BRCA1-deficient cells, where olaparib toxicity seems to arise from an inability to perform HRR even if NHEJ is inactivated (Bunting et al., 2010; Chen et al., 2017).

### ATM-deficient cells exhibit delayed DNA-end resection but do not accumulate unrepaired seDSBs

HRR of DSBs starts with DNA-end resection. While ATM is implicated in this process and is widely regarded as a HRR-promoting factor, its importance for HRR has been disputed, at least in the context of repairing DSBs arising from endonuclease activity (Kass et al., 2013; Rass et al., 2013). To explore ATM involvement in HRR of seDSBs, we treated cells with topotecan and assessed accumulation of RPA into nuclear foci, a well-established resection marker (Sartori et al., 2007). Notably *Atm*^*-/-*^ cells displayed reduced intensity of RPA foci 1 hour after continuous topotecan treatment, indicative of delayed DNA-end resection (**Fig. 5A; Fig. S4A**). Importantly, this was not due to slower replication or generation of less DNA damage in ATM-deficient cells, as co-staining with antibodies against the DNA damage marker Ser-139 phosphorylated histone H2AX (γH2AX), or measurement of replication, did not reveal any significant differences between WT and *Atm*^*-/-*^ cells (**Fig. 5A; Fig. S4A-C**). However, while RPA-focus intensity decreased after topotecan withdrawal in WT cells (reflecting RPA replacement with RAD51 and ensuing HRR) RPA focus intensity increased in *Atm*^*-/-*^ cells (**Fig. 5B**). Thus, the overall levels of RPA-focus formation/intensity during the experiment were very similar in cells containing or lacking ATM (**Fig. 5A-C; Fig. S4A).**

**Figure 5.**
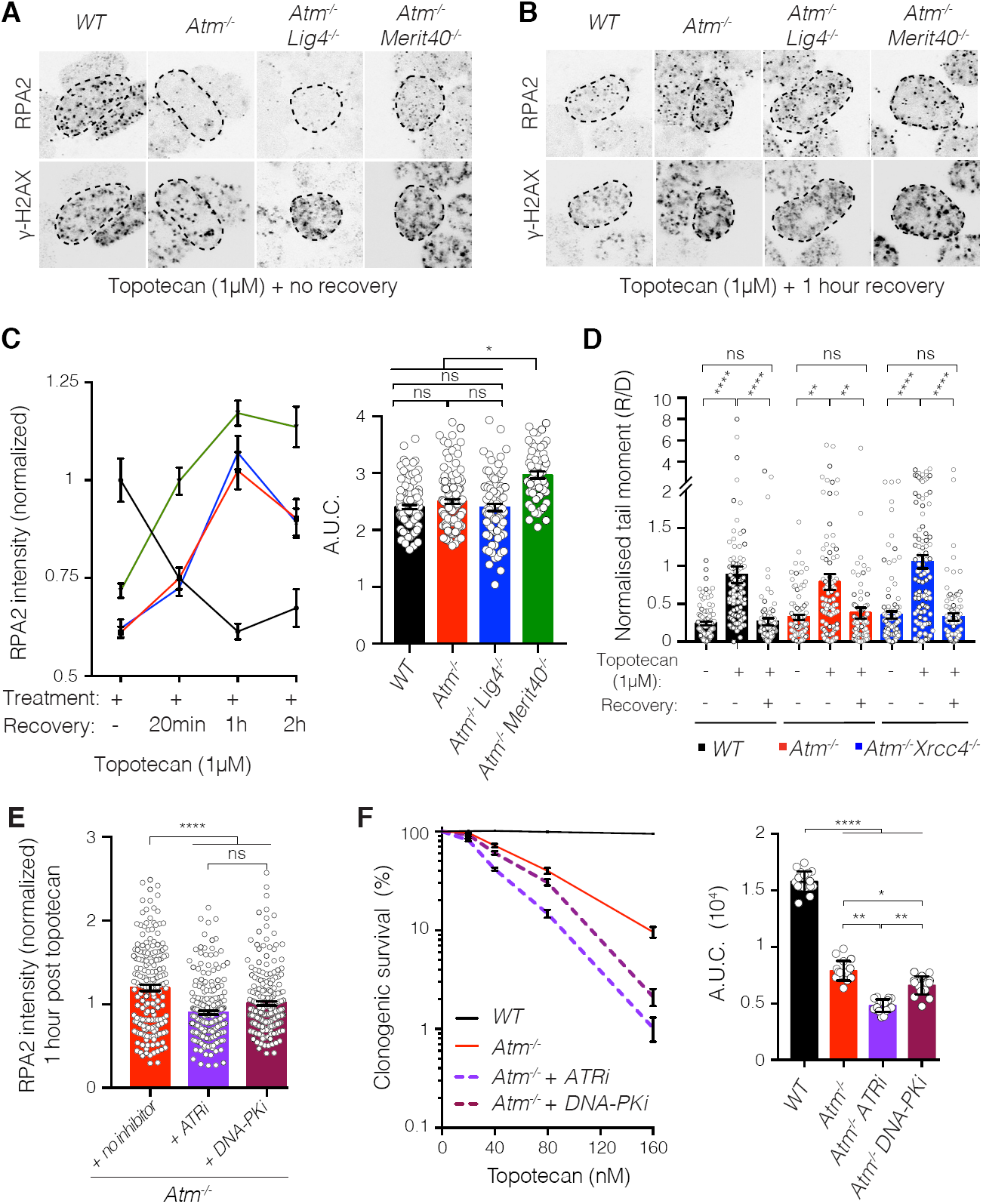
ATM-deficient cells can execute resection and do not accumulate unrepaired seDSBs upon topotecan treatment. **(A** and **B)** Representative images used for quantification of RPA2 foci accumulation in γH2AX-positive nuclei. *Atm*^*-/-*^*Lig4*^*-*^*/-* and *Atm*^*-/-*^*Merit40*^*-/-*^ cells are compared to WT and *Atm*^*-/-*^ cells upon topotecan treatment for 1h **(A)** or 1h after topotecan was removed **(B)**. For better visualization, color images were converted to black and white; representative color images are presented in **Figure S4**. Dashed outline indicates periphery of nuclei based on DAPI staining. **(C)** Quantification of topotecan-induced RPA2 foci formation, showing an initial delay in seDSB end-resection in *Atm*^*-/-*^ (n=122) and *Atm*^*-/-*^*Lig4*^*-/-*^ (n=77) cells compared to WT (n=110) cells (no recovery time point), but a recovery in resection during the two-hour timeframe after topotecan withdrawal. Overall, *Atm*^*-/-*^*Merit40*^*-/-*^ cells (n=57) show significantly higher resection when compared to all the other genotypes. RPA2 intensity quantifications were analyzed exclusively in #x03B3;H2AX-positive nuclei, representing S-phase cells that had encountered topotecan-induced TOP1-DNA cleavage complexes during replication. Cells were treated for 1h with 1µM topotecan and recovered for 20 minutes (20min), 1h or 2h without the drug. Graphs quantifying RPA intensity (left) and AUC (right) were generated by using GraphPad Prism 7. Bars represent mean ± SEM; ****p<0.0001; ns= not significant (p>0.05); two-tailed Student’s t test following F test to confirm equal variance; df=4. Data from n=3 individual experiments. **(D)** Neutral comet assays showing similar DNA damage generation and repair patterns upon seDSB induction in WT (n=86), *Atm*^*-/-*^ (n=80) and *Atm*^*-/-*^*Xrcc4*^*-/-*^ (n=89) cells. Cells were treated for 1h with 1µM topotecan, the drug removed and cells allowed to recover for a further 6h. Bar graphs represent the mean ± SEM of the normalized ratio of recovery to damage (R/D) tail moments. The graph was generated by using GraphPad Prism 7; ****p<0.0001; **p<0.01; ns= not significant (p>0.05); two-tailed Student’s t test following F test to confirm equal variance; df=4. Data from n=3 individual experiments. **(E)** Bar graph of the extent of RPA2 foci formation in ATM-deficient cells after 1 h recovery from a 1 h topotecan treatment (no inhibitor) compared to treatment with topotecan and ATR inhibitor (ATRi; AZD6738; 1µM) or DNA-PK inhibitor (DNA PKi; NU7441; 3µM). The inhibitors were added 1h before topotecan treatment, and samples were collected at the indicated time point. Bars represent mean ± SEM; *p<0.05; ns= not significant (p>0.05); two-tailed Student’s t test following F test to confirm equal variance; df=4. For each experiment data is pooled from n=3 individual experiments; n≥30 replicates in each time point. **(F)** Quantification of clonogenic survival showing significant increased sensitivity of *Atm*^*-/-*^ cells treated with ATR inhibitor or DNA PK inhibitor compared to *Atm*^*-/-*^ cells (no inhibitor) upon topotecan treatment; n=18/genotype.

To investigate whether ATR or DNA-PKcs could compensate for ATM’s role in promoting RPA loading, we treated cells with topotecan in the absence or presence of selective ATR or DNA-PKcs inhibitors. Inhibiting ATR, and to a lesser extent DNA-PKcs, reduced topotecan-induced RPA focus formation in ATM-deficient cells (**Fig. 5E**) and further increased their sensitivity to topotecan (**Fig. 5F**). These findings suggest that ATR and DNA-PKcs may cooperate with ATM at seDSB sites.

Collectively, the above findings implied that ATM deficiency does not prevent DNA resection but instead delays its kinetics, supporting our findings that showed no significant differences in the generation and repair of seDSBs produced by topotecan between WT and ATM-deficient cells, as assessed by neutral comet assays (**Fig. 5D**). Furthermore, loss of LIG4 or XRCC4 in *Atm*^*-/-*^ cells had no perceptible impact on topotecan-induced RPA focus generation or seDSB repair, implying that these NHEJ factors do not markedly affect resection in ATM-deficient settings (**Fig. 5C, D; Fig. S4A**). By contrast, *Atm*^*-/-*^*Merit40*^*-/-*^ cells exhibited overall higher levels of RPA focus intensity than *Atm*^*-/-*^ or *Atm*^*-/-*^*Lig4*^*-/-*^ cells (**Fig. 5A-C; Fig. S4A**), a finding in line with the documented role of the BRCA1-A in suppressing resection (Coleman and Greenberg, 2011; Hu et al., 2011).

### ATM-deficient cells mediate HRR of seDSBs arising at broken replication forks

Recent studies have documented persistent Ku foci at sites of seDSBs in cells treated with ATMi, suggesting that the hypersensitivity of such cells to TOP1 poisons reflects defective Ku removal impairing HRR of replication-associated seDSBs (Britton et al., 2013; Chanut et al., 2016b). However, although ATM-deficient cells displayed increased topotecan-induced Ku foci compared to WT, LIG4 depletion only partially reduced their numbers (**Fig. 6A, B**) despite LIG4 loss almost fully alleviating the topotecan hypersensitivity of ATM-null cells (see **Figs. 2** and **3**). As LIG4 catalytic activity drives topotecan toxicity in ATM-deficient cells (**Fig. 3B**), these results suggested that in such settings, Ku driven topotecan toxicity most likely reflects it promoting LIG4/XRCC4 recruitment and LIG4 catalytic activity at seDSBs. In this regard, we noted that ATM inhibition also increased XRCC4 focus formation following topotecan treatment (**Fig. S5A**). In accord with these observations and noting that ATM-dependent phosphorylation of CTIP counteracts Ku at seDSBs, CTIP depletion did not further enhance the sensitivity of ATM-deficient cells to topotecan (**Fig. 6C, D**), thus placing ATM and CTIP in an epistatic relationship. Furthermore, while CTIP depletion enhanced WT cell sensitivity to topotecan, it did not affect topotecan sensitivity in *Atm*^*-/-*^*Lig4*^*-/-*^ cells (**Fig. 6E, F**).

**Figure 6.**
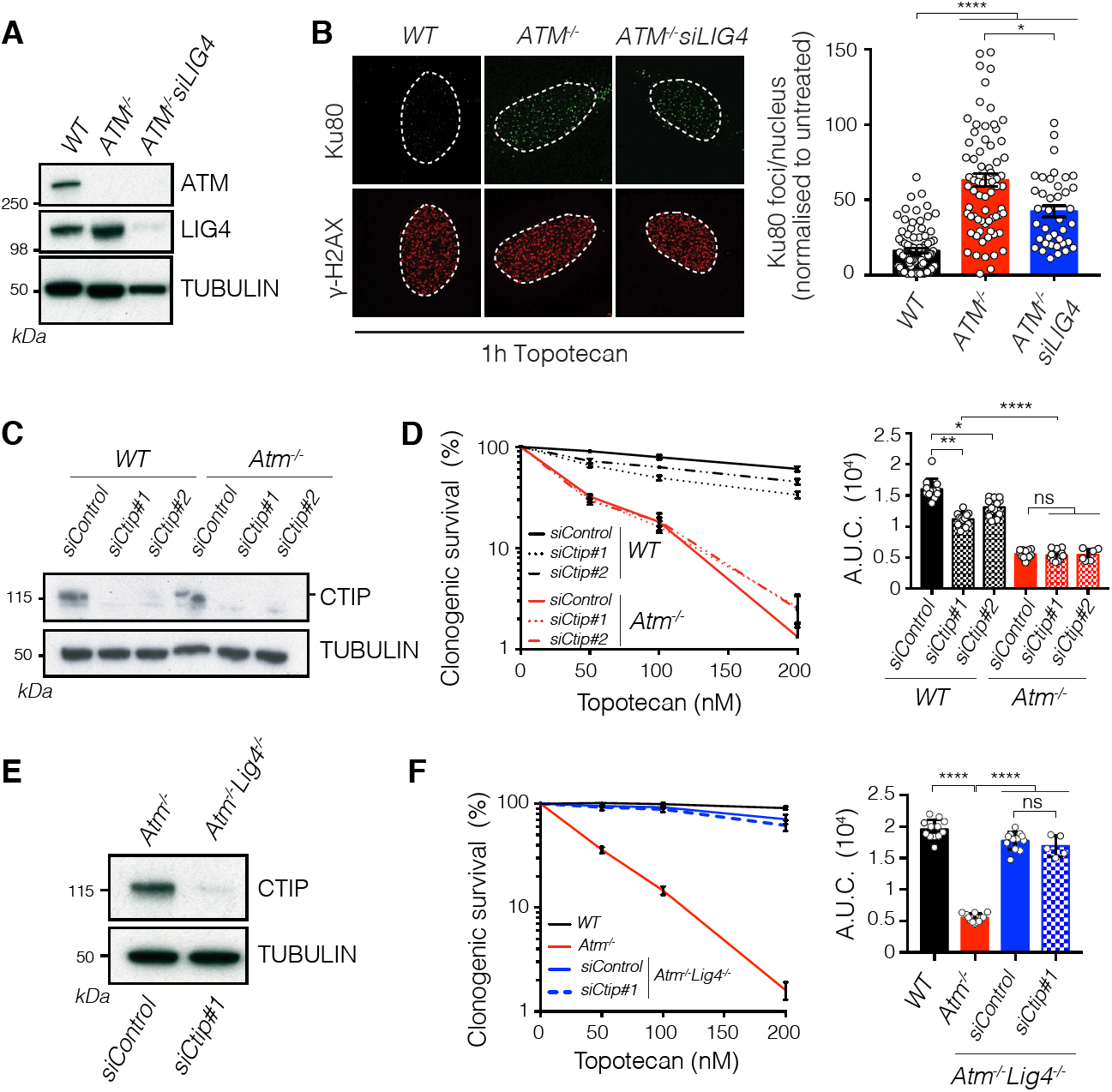
Impaired CTIP-dependent Ku removal plays a minor role in generating toxicity to topotecan in ATM-deficient cells. (**A**) Representative western blot showing absence of ATM protein in *ATM*^*-/-*^ U2OS cells. LIG4 proteins levels were analyzed in *ATM*^*-/-*^ cells treated with *siLIG4*. (**B**) Representative images and quantifications of Ku80 foci in #x03B3;H2AX-positive nuclei in U2OS cells treated with topotecan. Dashed outline indicates periphery of nuclei. Bars represent mean ± SEM; ****p<0.0001; *p<0.05; two-tailed Student’s t test following F test to confirm equal variance; df=4. Data from n=3 individual experiments. (**C**) Representative western blot on the representative genotypes upon transfection with siControl as compared to *siCtip #1* and *siCtip #2* and analyzed for CTIP protein levels.(**D**) Panels containing clonogenic survival (left) and AUC (right) upon topotecan treatment in cells depleted of *Ctip* in WT (*Atm*^*+/+*^; n=12 for each siRNA) and *Atm*^*-/-*^ (n=12 for each siRNA) cells as compared to control treatment with siControl (n=12/genotype). (**E**) Representative western blot analysis on the representative genotypes upon transfection with siControl as compared to *siCtip #1* and analyzed for CTIP protein levels. (**F**) Panels containing clonogenic survival (left) and AUC (right) upon topotecan treatment in cells depleted of *Ctip* in *Atm*^*-/-*^ *Lig4*^*-/-*^ (*Atm*^*-/-*^ *Lig4*^*-/-*^; n=12 for each siRNA) as compared to control treatment with siControl (n=12/genotype) in *Atm*^*+/+*^, *Atm*^*-/-*^ and *Atm*^*-/-*^ *Lig4*^*-/*^ backgrounds^*-*^. In panels (**D**) and (**F**) containing clonogenic survival curves (left) and AUC (right) were generated using GraphPad Prism 7. Bars represent mean ± SEM; ****p<0.0001;***p<0.001; **p<0.01; *p<0.05; ns= not significant (p>0.05); two-tailed Student’s t test following F test to confirm equal variance; df=4. Data from n=3 individual experiments.

Restart of a broken replication fork that has generated a seDSB requires HRR using the homologous sister chromatid as template. This mechanism involves RAD51-dependent strand invasion and formation of a Holliday junction, which upon resolution results in a SCE (Helleday, 2003). It has been shown that absence of CTIP or impaired MRE11 exonuclease activity results in defective RAD51 accumulation on seDSBs, due to persistence of Ku foci (Chanut et al., 2016b), thus implying that ATM-deficient cells would show defective SCE formation upon seDSB induction. Strikingly, we found that ATM deficiency did not affect topotecan-induced SCE formation when compared to WT cells or to *Atm*^*-/-*^*Xrcc4*^*-/-*^ or *Atm*^*-/-*^*Merit40*^*-/-*^ cells (**Fig. 7A, B;** for controls showing equivalent cell-cycle progression see **Fig. S4B, C**; for karyotypes and SCEs, see **Fig. S5B, C, S6A**). These results thereby supported our other findings indicating that, although delayed, HRR of seDSBs at broken replication forks can be completed in the absence of ATM, and that overall HRR efficiency is not overtly affected by LIG4/XRCC4 or components of the BRCA1-A complex. Furthermore, they suggested that persistence of Ku at seDSBs in the absence of ATM activity does not impair their resolution by HRR.

**Figure 7.**
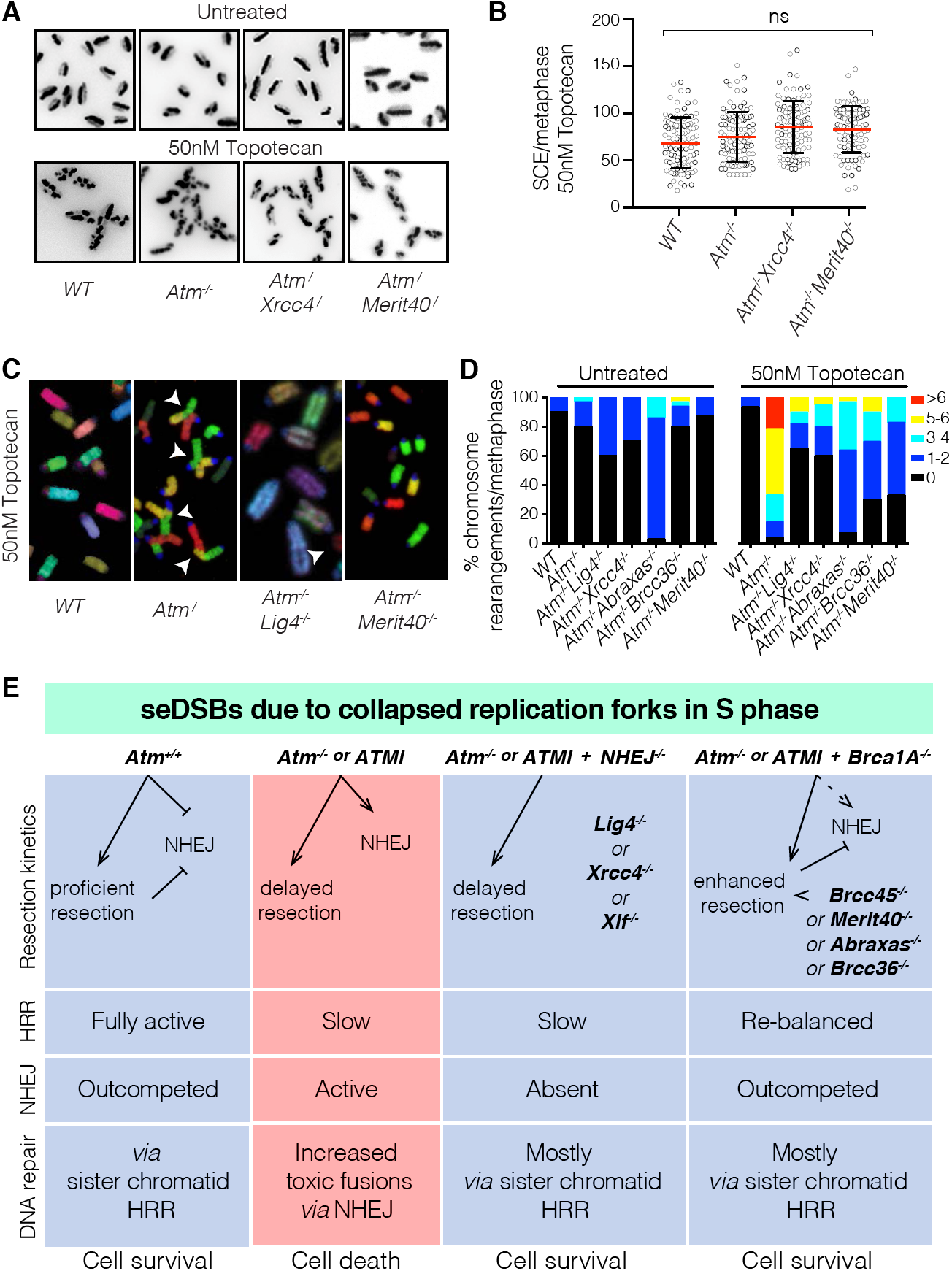
ATM counteracts toxic NHEJ of seDSBs in S phase. (**A** and **B)** ATM is not required for BIR-mediated repair of collapsed replication forks. Representative images **(A)** and quantification **(B)** of sister chromatid exchanges (SCEs) in cells treated with 50nM topotecan (n=100/genotype). Quantifications of chromosome numbers and SCEs in untreated cells are presented in **Figure S5C, D**. Scatter dot plots showing mean ± SD number of SCEs across the representative genotypes. Data from n=3 individual experiments. The graph was generated by using GraphPad Prism 7; ns= not significant (p>0.05); two-tailed Student’s t test following F test to confirm equal variance; df=4**. (C)** ATM is required to prevent toxic fusions upon formation of seDSBs. Representative images of metaphase spreads depicting multicolor fluorescent in-situ hybridization (M-FISH) using mouse 21-color painting chromosome probes. White arrows indicate fusions. Representative karyotypes are presented in **Figure S6A. (D)** Contingency graphs showing the percentages of chromosome rearrangements from chromosomal spreads of untreated cells and cells treated with 50nM topotecan, generated by using GraphPad Prism 7. n=3 individual experiments measuring n≥20 metaphases/genotype in each experiment were karyotyped. Statistical analysis is presented in **Figure S6C. (E)** Model for the role of ATM in the repair of seDSBs resulting from collapsed replication forks in S-phase of the cell cycle. Column 1, ATM promotes resection of seDSBs, thereby speeding up their repair by HRR and minimizing the time-window during which toxic NHEJ might take place. As shown, ATM also counteracts NHEJ by other mechanisms (see main text). Column 2, in the absence of ATM, seDSB resection is delayed and NHEJ is not suppressed, leading to some seDSBs being subject to illegitimate NHEJ, causing chromosome fusions and ensuing cell death. Column 3, inactivating NHEJ alleviates the hypersensitivity of ATM-null cells to agents that generate seDSBs because toxic, illegitimate NHEJ is absent. Column 4, modifying seDSB end-resection dynamics by loss of BRCA1-A complex components alleviates (rebalances) the seDSB resection defect of ATM-deficient cells, thereby minimizing the potential for illegitimate NHEJ.

### ATM-deficient cells accumulate toxic NHEJ-mediated chromosome aberrations

Significantly, we observed that while ATM-deficient cells were competent to eventually execute HRR of seDSBs, *Atm*^*-/-*^ cells treated with topotecan displayed chromosomal aberrations, especially involving fusions of chromatids from different chromosomes (**Fig. 7C, D; Fig. S6A, B**). Indeed, continuous exposure to topotecan resulted in almost all metaphases from *Atm*^*-/-*^ cells exhibiting at least one chromosomal aberration, while this was only seen in ∼5% of WT cells (**Fig. 7D**, right panel). Crucially, the extent of such chromosomal aberrations in *Atm*^*-/-*^ cells was markedly reduced in *Atm*^*-/-*^*Lig4*^*-/-*^ or *Atm*^*-/-*^*Xrcc4*^*-/-*^ double mutant backgrounds, and was also substantially decreased in *Atm*^*-/-*^*Abraxas*^*-/-*^, *Atm*^*-/-*^*Brcc36*^*-/-*^ or *Atm*^*-/-*^*Merit40*^*-/-*^ cells **(Fig. 7C, D; Fig. S6C)** paralleling the effects we observed on cell survival. Together, these findings supported a model in which topotecan-induced killing of ATM-deficient cells is largely mediated *via* the formation of chromosomal aberrations by a mechanism(s) that can be circumvented by inactivation of certain NHEJ components or deficiency in proteins of the BRCA1-A complex.

## DISCUSSION

*ATM* mutations are found in various cancers (Lawrence et al., 2014) and also cause the neurodegenerative and cancer-predisposition syndrome, Ataxia Telangiectasia (Shiloh and Ziv, 2013). Similar to loss-of-function of breast cancer susceptibility genes *BRCA1* or *BRCA2*, *ATM* loss or mutation causes hypersensitivity to various clinical DNA damaging agents (Holohan et al., 2013). Because ATM affects DSB resection, a key early step in HRR, hypersensitivity of ATM-deficient cancer cells to PARP inhibitors, TOP1 poisons and other S-phase DNA-damaging agents may arise from HRD (Lord and Ashworth, 2016).

Through CRISPR-Cas9 genetic screening and ensuing studies, we established that inactivation of genes encoding a subset of classical NHEJ proteins (LIG4, XRCC4, XLF) or components of the BRCA1-A complex (BRCC45, BRCC36, ABRAXAS, MERIT40) alleviates toxicity exerted by TOP1 and PARP inhibitors on ATM-null cells or in cells with catalytically-inactive ATM. Furthermore, we found that toxicity in ATM-null cells is largely mediated by LIG4 catalytic activity, implying that it is not the recruitment or persistence of Ku and the NHEJ machinery at seDSBs *per se* but lack of NHEJ-mediated DNA-end ligation that drives drug sensitivity in ATM-deficient settings. Importantly, in accord with published data (Kass et al., 2013; White et al., 2010), we found that ATM-null cells, although presenting delayed resection kinetics, are proficient in HRR of replication-associated seDSBs. We thus conclude that *ATM*-mutant cells fail to prevent some seDSBs being converted into toxic chromosomal aberrations by NHEJ during S-phase, which ultimately kill cells harboring them. Accordingly, resistance to TOP1 or PARP inhibitors ensues in ATM-deficient cells when such toxic NHEJ is prevented; either directly by loss of NHEJ end-ligation factors, or indirectly *via* inactivating BRCA1-A complex components, modifying DSB resection dynamics to increase HRR efficiency (**Fig. 7E**).

The resistance mechanisms we describe have important implications for our understanding of seDSB repair. First, our results show that the overall efficiency of HRR of seDSBs is not strongly affected by ATM loss. Second, in the absence of ATM, CTIP is not required for HRR of seDSBs, as its depletion does not further increase topotecan sensitivity. Third, we observed that HRR of seDSBs as measured by SCEs is not impaired by the absence of ATM, even though Ku foci persist under these circumstances. Fourth, we established that absence of LIG4 is sufficient to suppress the sensitivity of ATM-deficient cells to topotecan, even though it has a minor impact on Ku persistence at seDSBs. Collectively, these results imply that toxicity to seDSB-inducing agents in ATM-deficient cells is primarily driven by the completion of the ligation step of NHEJ at a limited number of seDSBs, and not by the inability to load RAD51 onto resected DSBs due to inefficient Ku removal. As ATM phosphorylates hundreds of substrates (Matsuoka et al., 2007), it may operate at multiple, other levels (Chanut et al., 2016a) to prevent NHEJ and ensure HRR of broken replication forks. Indeed, by carrying out SILAC mass-spectrometry analysis to identify ATM-dependent phosphorylations upon TOP1 poisoning, we identified over 100 phosphorylation sites that are under ATM control (**Fig. S7A-D, Dataset S7**). These data argue that ATM orchestrates responses to seDSB accumulation at multiple levels that could collectively control resection speed as well as inhibit NHEJ.

Notably, DNA-PKcs or PAXX deficiency does not rescue the hypersensitivity of ATM-deficient cells to TOP1 or PARP inhibitors, implying that toxic end-joining events at seDSBs can happen independently of these factors. This is in line with findings indicating that unlike XLF, PAXX does not play a significant role in NHEJ in S/G2 phase cells (Kumar et al., 2016). It is tempting to speculate that this might reflect replication-associated seDSBs being relatively simple (“clean”) substrates that can be ligated together by the actions of the Ku, XRCC4, XLF and LIG4 proteins without DNA-end processing activities dependent on DNA-PKcs (Reynolds et al., 2012).

Our findings also highlight how ATM-deficient cells’ hypersensitivity to seDSB-inducing agents is mechanistically different from the scenario in BRCA1-deficient cells, where it arises from an inability to perform HR even if NHEJ is inactivated (Bunting et al., 2010; Chen et al., 2017). Accordingly, the genetic suppression mechanisms that we have defined for *ATM*-mutant cells appear distinct from those reported in *BRCA1*-mutant contexts, where suppression occurs specifically upon inactivation of DNA-end resection and HR-antagonist protein 53BP1 and its interactors (Chen et al., 2017; Xu et al., 2015). However, we cannot exclude the possibility that 53BP1 recruitment to seDSBs is promoted by ATM function, as has been shown at double-ended DSBs (Zimmermann and de Lange, 2014). Consequently, we suggest caution when interpreting HRD in isolation as a clinical prognostic tool. The HRD score (Myriad Genetics HRD™) (Abkevich et al., 2012) or the HRDetect mutational signature model (Davies et al., 2017) have been proposed as predictive biomarkers of treatment response to agents such as PARP inhibitors, regardless of etiology or mechanism-of-action. Based on our findings, we suggest that these approaches could miss opportunities presented by deficiencies in ATM and/or in other factors involved in suppressing NHEJ at broken DNA-replication forks during S phase (Polak et al., 2017).

Intrinsic or acquired tumor cell resistance to established chemotherapeutics, and towards newer molecularly-targeted agents such as PARP inhibitors, is a major problem in cancer management (Kumar-Sinha and Chinnaiyan, 2018). Understanding the molecular bases for drug resistance is thus crucial to establish better patient stratification and combination chemotherapy regimens, as well as to better understand mechanisms and relationships between cellular DDR and other processes. Based on our findings, we suggest that exploring the genetic and transcriptional status of BRCA1-A complex members and NHEJ components in ATM-deficient tumors might help predict responses to seDSB-inducing agents such as TOP1 poisons or PARP inhibitors. Furthermore, the identification of deficiencies in the same genes as driving resistance to the combination of irinotecan and ATM inhibitors in human cancer cells provides further evidence for the potential translation of our findings to clinically-relevant scenarios. Finally, we note that our finding that LIG4 or XRCC4 loss further sensitizes ATM-deficient cells to IR highlights the potential for exploiting a resistance mechanism towards one drug-type as a vulnerability towards another therapeutic regime.

## ACKNOWLEDGEMENTS

We thank all S.P.J. laboratory members as well as Dr. Andrew Blackford and Dr. Sébastien Britton for discussions, and Kate Dry for editorial assistance. We thank Richard Collins for help with the mouse xenografts. Research in the SPJ laboratory is funded by Cancer Research UK (programme grant C6/A18796), Wellcome Trust Investigator Award (206388/Z/17/Z) and an Action for A-T grant (16GUR02). Institute core infrastructure funding is provided by Cancer Research UK (C6946/A24843) and the Wellcome Trust (WT203144). SPJ receives salary from the University of Cambridge. GB and JVF were funded by Cancer Research UK programme grants C6/A11224 and C6/A18796, and the Ataxia Telangiectasia Society. DP was funded by Cancer Research UK studentship C6/A21454. JC was funded by Cancer Research UK programme grants C6/A11224 and C6/A18796. The PB lab is supported by the Emmy Noether Program (BE 5342/1-1) from the German Research Foundation and the Marie Curie Career Integration Grant from the European Commission (630763).

## AUTHOR CONTRIBUTION

GB, JVF and SPJ wrote the manuscript. GB and JVF assembled the figures and supervised/were involved in all the experiments. DP performed and analyzed RPA resection assays, DNA comet assay, generated BRCA1-A double mutant cells and performed clonogenic survivals with help from JC. DP performed and quantified high-resolution microscopy. JC helped with clonogenic survival assays and cell line generation. MD and MSC generated human RPE1 mutant cells. FMM derived the ATM KO U2OS cells. AB quantified the SCEs and helped with the CRISPR screen. MW helped with the ESC derivation. BF and FY performed the FISH experiments and quantification. EC, MG and MH performed the human cancer cell lines analysis. CEO and TS harvested and analyzed the human cancer samples. KY, HP and AB helped with the CRISPR-Cas9 library preparation and data analysis. MO and PB performed and analyzed the SILAC-mass spectrometry experiments. EM helped with the CRISPR-Cas9 screen design, library preparations, sequencing and supervised the screen data analysis. GB performed the CRISPR-Cas9 screen and helped with and coordinated all the mouse work with help from DJA. All authors commented on the manuscript and figures. SPJ supervised the work.

